# Ancestral gene synteny reconstruction improves extant species scaffolding

**DOI:** 10.1101/023085

**Authors:** Yoann Anselmetti, Vincent Berry, Cedric Chauve, Annie Chateau, Eric Tannier, Sèverine Bérard

## Abstract

We exploit the methodological similarity between ancestral genome reconstruction and extant genome scaffolding. We present a method, called ART-DECO that constructs neighborhood relationships between genes or contigs, in both ancestral and extant genomes, in a phylogenetic context. It is able to handle dozens of complete genomes, including genes with complex histories, by using gene phylogenies reconciled with a species tree, that is, annotated with speciation, duplication and loss events. Reconstructed ancestral or extant synteny comes with a support computed from an exhaustive exploration of the solution space. We compare our method with a previously published one that follows the same goal on a small number of genomes with universal unicopy genes. Then we test it on the whole Ensembl database, by proposing partial ancestral genome structures, as well as a more complete scaffolding for many partially assembled genomes on 69 eukaryote species. We carefully analyze a couple of extant adjacencies proposed by our method, and show that they are indeed real links in the extant genomes, that were missing in the current assembly. On a reduced data set of 39 eutherian mammals, we estimate the precision and sensitivity of ART-DECO by simulating a fragmentation in some well assembled genomes, and measure how many adjacencies are recovered. We find a very high precision, while the sensitivity depends on the quality of the data and on the proximity of closely related genomes.

## Introduction

Knowledge of genome organization (gene content and order) and of its dynamics is an important question in several fields such as cancer genomics [1, 2, 3], to understand gene interactions involved in a common molecular pathway [4], or evolutionary biology, for example to establish a species phylogeny by comparative analysis of gene orders [5].

On one side, studying genome organization evolution, and in particular proposing gene orders for ancestral genomes, requires well assembled extant genomes, while, on the other side, the assembly of extant genomes can in return benefit from evolutionary studies. This calls for integrative methods for the joint scaffolding of extant genomes and reconstruction of ancestral genomes.

The reconstruction of ancestral genome organization is a classical computational biology problem [6], for which various methods have been developed [7, 8, 9, 10, 11, 12, 13]. The rapid accumulation of new genome sequences provides the opportunity to integrate a large number of genomes to reconstruct their structural evolution. However, a significant proportion of these genomes is incompletely assembled and remains at the state of contigs (permanent draft) as illustrated by statistics on GOLD [14]. To improve assemblies, methods known as scaffolding were developed to order contigs into scaffolds. Most scaffolding methods use either a reference genome, or the information provided by paired-end reads, or both [15, 16, 17, 18, 19, 20, 21, 22]. We refer to Hunt et al. [23] for a detailed comparative analysis of recent scaffolding methods.

In recent developments, scaffolding methods taking into account multiple reference genomes and their phylogeny have been proposed [24, 25, 26, 27]. It suggests a methodological link with ancestral genome reconstructions: if ancestral genes are considered as contigs, scaffolding extant or ancestral contigs in a phylogenetic context differs only in the location of the considered genome within the phylogeny (leaf or internal node). This link has been observed [28] and exploited to develop a method for combining scaffolding and ancestral genomes reconstruction [29]. However, the latter, due to the computational complexity of the ancestral genome reconstruction problem, is currently limited to a few genomes and to single-copy universal genes.

We propose to overcome this computational complexity by considering independent ancestral gene neighborhood reconstructions [12] instead of whole genomes, which allows to scale up to dozens of whole genomes and to use as input data genes with complex histories. We develop a method that scaffolds ancestral and extant genomes at the same time. The algorithm improves over previous methods of scaffolding by the full integration of the inference of evolutionary events within a phylogenetic context.

The principle of our method is imported from DECO [12]. So we call it ART-DECO for Assembly Recovery through DECO. DECO is an algorithm for ancestral synteny reconstruction. It is a dynamic programming scheme on pairs of reconciled gene tree, generalizing the classic dynamic programming scheme for parsimonious ancestral character reconstructions along a tree. It computes a parsimonious set of ancestral gene neighborhoods, the cost being computed as the weighted sum of gains and losses of such neighborhoods, due to genome rearrangements. In addition to DECO, ART-DECO considers a linkage probability for each couple of genes in extant genomes, that is included in the cost function in order to be able to propose gene neighborhoods in extant as well as in ancestral genomes.

We implemented ART-DECO and tested it on several data sets. First we reproduced the experiment of [29] on 7 tetrapod genomes limited to universal unicopy genes, with comparable accuracy. Then we used all genes from 69 eukaryotic genomes from the Ensembl database [30]. The program runs in about 18h and is able to propose ancestral genome structures and thousands of extant scaffolding linkages. We examine in details one of them, chosen randomly on the panda genome, and show why it seems reasonable to propose it as an actual scaffolding adjacency. Then on a reduced data set of 39 whole mammalian genomes, we tested the precision and sensitivity of the scaffolding performed by ART-DECO by simulating artificial fragmentation of the human or horse genomes, removing up to 75% of the known gene neighborhoods of these well assembled genomes, and comparing the removed adjacencies with the ones proposed by ART-DECO. We measure a *>* 95% precision, while sensitivity, as expected, depends on the quality of the data and on the presence of closely related extant genomes. This denotes the domain of efficiency of our method: a vast majority of proposed adjacencies can be considered with confidence, but the final resulting scaffolding is still incomplete.

## Ancestral and extant adjacencies

We describe the ART-DECO algorithm for the joint reconstruction of ancestral genomes and scaffolding of extant genomes. An overview is depicted in Figure 1.

**Figure 1.**
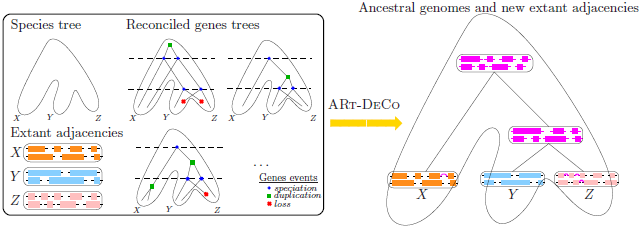
Input and output of the ART-DECO method. The left box shows the input of ART-DECO: a species tree (here on extant species *X*, *Y* and *Z*), the adjacencies in the genome of extant species (each colored block represents a contig, that is, a linear arrangement of genes, linked by adjacencies) and the reconciled genes trees for their genes. The output of ART-DECO is shown on the right-hand side in magenta color: the method computes both new adjacencies for extant species and contigs for the ancestral species.

### Input

The input of the method is

- A species tree with all considered genomes and their descent pattern; We suppose the number of chromosomes of each extant genome is known.
- A set of genes for all considered genomes, clustered into homologous families; for each family a rooted gene tree depicts the descent pattern of the genes.
- A set of *adjacencies*, *i.e.*, pairwise relations between neighboring genes *AB* on extant chromosomes. Genes *A* and *B* are called the *extremities* of the adjacency *AB*. We consider as neighbors two genes that are not separated by another gene present in the dataset, but a relaxed definition can be used with no impact on the method itself.

Internal nodes of the species tree are labeled with ancestral species (we always consider ancestral species at the moment of a speciation) and leaves are labeled with extant species. Gene trees are reconciled with the species tree: all ancestral genes are labeled by the ancestral species they belong to, so the input yields a gene content for all ancestral species. Genes and species are partially ordered by the descent relation, so we may speak of a last, or lowest, or most recent common ancestor. Here, as in [12], we use a reconciliation minimizing the number of duplications and losses of genes.

A module of ART-DECO is able to produce a suitable input from the raw Ensembl Compara [30] gene tree files and a species tree if needed. Once the input is given, two preliminaries are necessary: partitioning extant adjacencies and computing an a priori adjacency probability for each extant species. They are detailed in the two following subsections.

### A partition of extant adjacencies

The goal of this step is, without loss of generality, to reduce the analysis of the whole data set to the independent analysis of pairs of gene trees and adjacencies, each having an extremity in each of the gene trees. Moreover, we want that the roots of the two gene trees correspond to ancestral genes mapping to the same ancestral species.

The partition is realized thanks to a necessary condition for two adjacencies to share a common ancestor. Two adjacencies *A*_1_*B*_1_ and *A*_2_*B*_2_, for genes *A*_1_, *A*_2_, *B*_1_, *B*_2_, may have a common ancestor *AB* only if *A*_1_ and *A*_2_ (respectively *B*_1_ and *B*_2_) are in the same gene family, so have a common ancestor *A* (respectively *B*), and such that *A* and *B* belong to the same species. In other words, the ancestral adjacency has the possibility to exist only when the genes of this adjacency are present in a same ancestral species.

It is easy to check that this relation is an equivalence relation, which then partitions adjacencies into equivalence classes. Each equivalence class *C* can be represented by two ancestral genes: they are the most ancient distinct *A* and *B* genes involved in the two-by-two comparisons of adjacencies *A*_1_*B*_1_ and *A*_2_*B*_2_ in this class. Necessarily every adjacency in this class has a gene which is a descendant of *A*, and another which is a descendant of *B*. *A* and *B* are in the same species (ancestral or extant), and cannot be the descendant one of another.

For a node *N* of a gene tree *T*, *T*(*N*) is the subtree of *T* rooted at *N*. Consider the two disjoint subtrees *T*(*A*) and *T*(*B*). All adjacencies in the equivalence class *C* have one extremity which is a leaf of *T*(*A*) and the other which is a leaf of *T*(*B*). So each equivalence class may be treated independently from the other, and the input can be restricted, without loss of generality, to *T*(*A*) and *T*(*B*).

### An *a priori* probability for all adjacencies

Given two extant genes *v*_1_ and *v*_2_ from the same extant genome *G*, we give an *a priori* probability that there is an adjacency between *v*_1_ and *v*_2_. If the genome *G* is perfectly assembled, then this probability is given by the input, that is, it is 1 if there is an adjacency in the input and 0 otherwise. But if the genome *G* is not perfectly assembled, then this probability depends on the quality of the assembly. It will allow the program to propose more adjacencies in extant genomes that are more fragmented.

We note *n* the number of contigs in an extant genome (which is the number of genes minus the number of adjacencies if all contigs are linear arrangements of genes), and *p* the number of chromosomes. We always have *n* ≥ *p* > 0. All contigs are assumed to have two distinct extremities.

We call a *solution* of the scaffolding problem a set of *n − p* adjacencies between the extremities of contigs, which forms *p* chromosomes from the *n* contigs. The number of different solutions for given *n* and *p* is denoted by *f*(*n, p*).

Let *v*_1_ and *v*_2_ be extremities of two different contigs; the *a priori* probability *P*(*v*_1_ ∼ *v*_2_) of *v*_1_ and *v*_2_ to be adjacent if they are not seen adjacent in the data and *n* > *p* is:

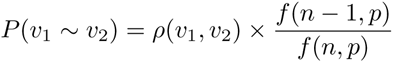

where *ρ* is a correction function which is equal to 4 if *v*_1_ and *v*_2_ are the only genes in their contigs, 2 if one of *v*_1_ *v*_2_ is the only gene in its contig but not the other, and 1 otherwise. If *n* = *p*, we have *P*(*v*_1_ *∼ v*_2_) = 0 if the adjacency *v*_1_*v*_2_ is not in the data, and *P*(*v*_1_ *∼ v*_2_) = 1 otherwise.

For the computation of *P*(*v*_1_ *∼ v*_2_) we use the following formula for *f*(*n, p*).

#### Lemma 1

*For each n ≥* 1 *and p* ∈ ℕ*, *we have:*

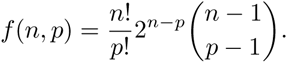

*Proof* First remark that the formula *f*(*n, p*) can be extended to the case where *p > n* and to the case where *n ≥* 1 and *p* = 0, by setting its value to 0 in those cases (there is no possible way to transform *n* contigs into *p* chromosomes). In those cases, the equality is still true, since 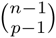 is then equal to 0. Thus, in all what follows, we use this extension of definition when needed.

We proceed now by induction on *n ≥* 1.

Base case: *n* = 1, it is straightforward since 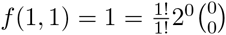, and for *p >* 1, we have 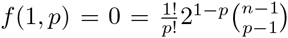, since the binomial coefficient is equal to 0 in this case.

Induction: we suppose that for each *k ≤ n*, for each *p* ∈ ℕ*\**, we have:

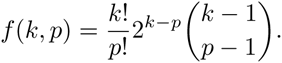

We consider *f*(*n* + 1, *p*), for a fixed *p* ∈ ℕ*. We sum over all possibilities for one specific chromosome to be composed of *x* contigs. This gives the recurrence formula

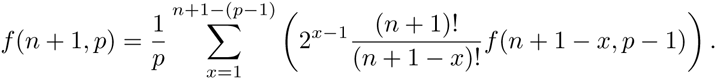

Where 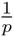 is used to avoid couting the same solution several times. 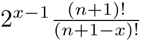 can be written 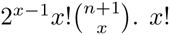 representing the number of possibilities to sort *x* contigs, 2^*x−*1^ allows to take into account contig orientations and 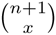 representsthe number of possibilities to pull *x* contigs of *n* + 1.

By induction hypothesis, we have:

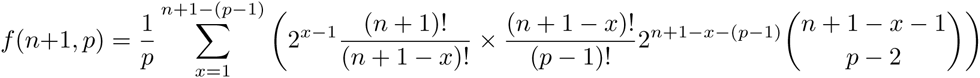

which simplifies into:

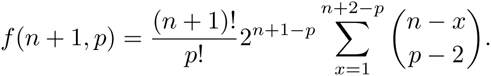

We change the variable in the sum, let *h* = *n − x*. Then, we have:

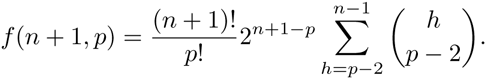

By the Hockey-stick’s identity, namely for all 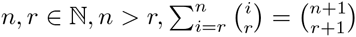 we finally obtain:

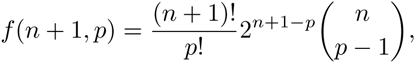

which concludes the proof.

The expression of *f*(*n, p*) leads to the following simple expression for *P*:

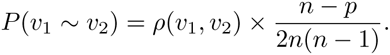

We define 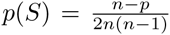 the part of this formula that does not depend on *v*_1_ and *v*_2_, as an assembly fragmentation measure for genome *S*.

### A Dynamic programming scheme

We largely refer to DECO [12] for a full description of the dynamic programming scheme, and only describe the overall principle and the differences we introduce. Adjacencies are constructed between ancestral genes (equivalently internal gene tree nodes), and propagate along gene trees. For two nodes *v*_1_ and *v*_2_ defining genes belonging to the same (ancestral or extant) species, we define a *solution* as a descent pattern of ancestral and extant adjacencies explaining the input extant adjacencies that have an extremity in *T*(*v*_1_) and another in *T*(*v*_2_). So a solution is a set of ancestral adjacencies and descent relations linking ancestral and extant adjacencies. The cost of a solution is the cumulative cost of gains and breakages of adjacencies (due to rearrangements) in the descent pattern, according to an individual cost for gains (*Gain*) and breakages (*Br*).

More precisely we define two costs *c*_0_(*v*_1_, *v*_2_) (respectively *c*_1_(*v*_1_, *v*_2_)), which are the minimum cost previously mentioned, given that there is an (respectively there is no) adjacency between *v*_1_ and *v*_2_ in a solution. All *c*_0_ and *c*_1_, for every couple *v*_1_ and *v*_2_, can be computed by the dynamic programming scheme described in [12]. ART-DECO and DECO have the same time complexity, that is *O*(*g*^2^ × *k*^2^) where *g* is the number of gene trees in the input and *k* be the maximum size of a tree.

The main difference is that in [12] extant genomes were supposed to be perfectly assembled and in particular, if *v*_1_ and *v*_2_ are extant genes (or equivalently gene tree leaves, which corresponds to Case 1 in [12]), then DECO would use the following scoring rules:

*c*_0_(*v*_1_, *v*_2_) = *∞* and *c*_1_(*v*_1_, *v*_2_) = 0 if *v*_1_*v*_2_ is an adjacency in the data, otherwise *c*_0_(*v*_1_, *v*_2_) = 0 and *c*_1_(*v*_1_, *v*_2_) = *∞*.

Here we modify these rules (it is the only case different from the dynamic programming equations given in [12] and Additional file 1) and propose instead that

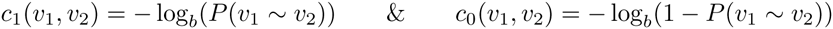

These formulas define a cost system which is consistent with the previous one: when *n* = *p* (perfectly assembled genomes) it gives the same result. When it is not the case, the costs are between 0 and *∞* as the probabilities go from 0 to 1.

We left the basis of the logarithm as a variable *b*. Giving a value to *b* determines a sensitivity for finding new adjacencies. It can be dependent on the genome *S* hosting *v*_1_ and *v*_2_. We choose the basis so that *c*_1_(*v*_1_, *v*_2_) *< c*_0_(*v*_1_, *v*_2_) + *Br* where *Br* is the cost of an adjacency breakage. Thus an adjacency is systematically proposed when it is inferred in the closest ancestor of *S*. The adjacency is obviously not always true in that case, because a rearrangement can have broken it in *S*. But it is a necessary condition to be able to propose any adjacency. If a genome is highly fragmented, proposing such an adjacency is more likely to lead to a true scaffolding adjacency than to cancel an evolutionary rearrangement. The relation *c*_1_(*v*_1_, *v*_2_) *< c*_0_(*v*_1_, *v*_2_) + *Br* yields

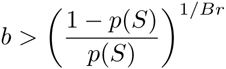

where *p*(*S*) represents the fragmentation of the genomes hosting *v*_1_ and *v*_2_, defined in the previous section. Preliminary experiments show that there is indeed a phase change in the number of inferred adjacencies when *b* reaches the right hand side of the above equation (Figure 2). Above this value, the number of inferred adjacencies is mainly constant, while it changes radically for smaller values. In following experiments we then fixed *b* to 1.05 times the right hand side of the above equation, in order to be sure to be on the plateau following the phase change.

**Figure 2.**
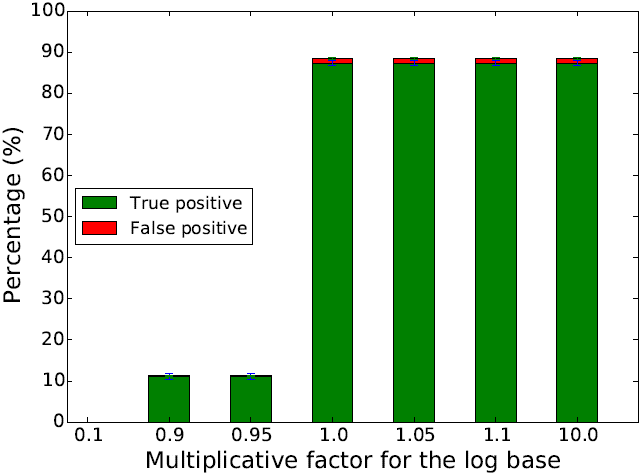
Determination of a good value for base log b. We simulated 550 fissions on a data set of 7 tetrapod species (see Section Results) and evaluate the ability of ART-DECO to recover broken adjacencies by the simulated fissions for different values for the base log *b*. On the *x* axis is the multiplicative factor of 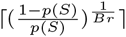, where *Br* = 1. As we can see on the graph, there is a phase change at 1.0, meaning that from this value a good number of adjacencies can be proposed. Increasing the multiplicative factor does not qualitatively change the results. This experiment was repeated for different numbers of simulated fissions (see Additional file 2) and different species trees, and in all cases results exhibited the same profile.

### Exploration of the solution space

The dynamic programming scheme of DECO allows the quantitative exploration of the whole solution space. This has been developed, in the DECO model, in [31], where it was shown how to explore all solutions (*i.e.* evolutionary histories for adjacencies) under a Boltzmann probability distribution defined as follows: for a given instance (pair of gene trees and set of extant adjacencies) with solution space *S*, the parsimony score of an adjacency history *h* is denoted by *s*(*h*), and the Boltzmann probability of *h* is defined as *e*^−*s*(*h*)/*kT*^/∑_*g∈S*_ *e*^−*s*(*g*)/*kT*^. Here *kT* is a constant that can be used to skew the probability distribution: when *kT* is small, parsimonious histories dominate the distribution, while a large *kT* leads to a more uniform distribution over the whole solution space.

This allows to associate to a feature of a solution (here an ancestral adjacency) a support defined as the ratio between the sum of the probabilities of the solutions that contain this feature and the sum of the probabilities of all solutions. This approach has been implemented in the DeClone software [31]. We integrated this possibility to ART-DECO and thus associate a support to both extant and ancestral adjacencies. Computations were run with a value of the *kT* constant equal to 0.1 to ensure that the Boltzmann distribution is highly dominated by optimal and slightly sub-optimal solutions. This value was chosen after preliminary tests on a subset of instances that showed that scenarios sampled with this value of *kT* were in very large majority optimal scenarios.

## Results

We tested ART-DECO on three data sets. The first one is composed of 7 tetrapod species with only universal unicopy genes, and aims at comparing our method with the method of Aganezov *et al.* [29]; on this data set, we obtain comparable results. Then we ran ART-DECO on the complete Ensembl Compara [30] database, including 69 eukaryotic species and 20,279 gene families with arbitrary numbers of duplications and losses. This shows that ART-DECO scales up and can process large data sets of whole genomes; for this data set we examine carefully one scaffolding adjacency proposed by ART-DECO in the poorly assembled panda genome and provide evidence it is likely a true scaffolding adjacency. The third data set we consider is the restriction to the 39 eutherian mammals genomes of the previous data set. The computational efficiency of ART-DECO allows to reproduce the computation many times with simulated missing adjacencies, and replicates to obtain empirical error bars on the measures. We performed all experiments with fixed costs for adjacency gains (*Gain*) and breakages (*Br*), respectively set to 3 and 1. There are several reasons for this discrepancy: first the actual number of adjacencies is very low compared to the space of possible adjacencies, which makes more probable to break a particular one 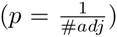 than to gain a particular one 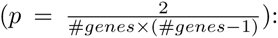 there is a huge unobserved space of possible solutions that should affect the costs; second it has been remarked that good statistical estimates of genomic distances when genomes are coded by the presence or absence of adjacencies are obtained with a state of possible adjacencies 3 times larger than the number of adjacencies (see paper “Moments of genomes evolution by Double Cut-and-Join”, BMC bioinformatics, to appear).

### Seven tetrapods - comparison with the method of Aganezov *et al.* [29]

By querying Biomart [32], we produced a data set similar to the one described in Aganezov *et al.* [29]: it consists in 8,818 universal unicopy gene families from Human, Chimp, Macaque, Mouse, Rat, Dog and Chicken. The latter was not present in the data set of Aganezov *et al.* [29], and we included it here because of a fundamental difference between the two methods: our method works with rooted phylogenies whereas Aganezov *et al.* [29] is not sensitive to the position of the root. This means that our method cannot scaffold an outgroup species, simply because, for any adjacency absent from the outgroup, it is more parsimonious to assume it is gained in all ingroup species. So we just added a distant outgroup to scaffold the 6 species used in [29].

We produced different sets of randomly fragmented genomes by considering *n* = 50 to *n* = 1050 random artificial breaks (or “fissions”) in genomes, sticking to the described experiments in [29]. This means we simply removed *n* random adjacencies per genome from the data set. For each *n*, we replicated the experiment 30 times.

For each replicate with *n* random artificial adjacency breaks, let *T P* be the number of removed adjacencies that ART-DECO retrieves and *F P* be the number adjacencies not in the removed ones but proposed by ART-DECO.

We measured, following [29], approximations of the sensitivity and precision:

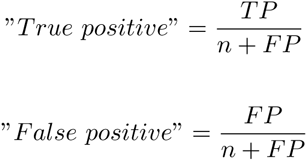

Aganezov *et al.* report that “*T rue positive*” takes values between 75% and 87%, and “*False positive*” takes values from 0.5% to 9%, varying in function of *n*. We report similar values for all our experiments (see Table 1).

**Table 1.** Statistics on adjacencies recover by ART-DECO on 7 tetrapods dataset with different number of simulated breaks.

| #Breaks (n) | 50 | 150 | 250 | 350 | 450 | 550 |
| --- | --- | --- | --- | --- | --- | --- |
| *TP* | 283 | 829 | 1364 | 1895 | 2418 | 2922 |
| *FP* | 14 | 16.5 | 21 | 24.5 | 32 | 40 |
| *True positive* | 88.64% | **89.98%** | 89.38% | 89.00% | 88.32% | 87.35% |
| *False positive* | 4.40% | 1.78% | 1.39% | **1.15%** | 1.18% | 1.19% |
| #Breaks (n) | 650 | 750 | 850 | 950 | 1050 |  |
| *TP* | 3431 | 3917 | 4398 | 4875 | 5338 |  |
| *FP* | 46 | 57 | 63 | 73.566 | 83 |  |
| *True positive* | 86.84% | 85.87% | 85.12% | 84.38% | 83.58% |  |
| *False positive* | 1.17% | 1.25% | 1.22% | 1.27% | 1.30% |  |

Thus, on small data sets and discarding gene families with complex histories, we obtain similar performance. The next experiments illustrate that the contribution of our method is then to be able to process much larger and much more complex data sets.

### 69 eukaryotes - a proof of scaling up

We ran ART-DECO on the full Ensembl Compara database (1,222,543 protein coding extant genes in 69 extant species) in about 18h. The input contains 1,023,492 adjacencies in the extant genomes, showing that many genomes assemblies are highly fragmented, from 11 chromosomes for the perfectly assembled opossum genome to 12,704 contigs for the wallaby genome, an order of magnitude comparable to the number of genes. In Figure 3, the black bars show the proportion of genes with 0, 1 or 2 syntenic neighbors in the extant input genomes. Around 30% of genes have at most one neighbor, while we would expect less than 1% for perfectly assembled genomes.

**Figure 3.**
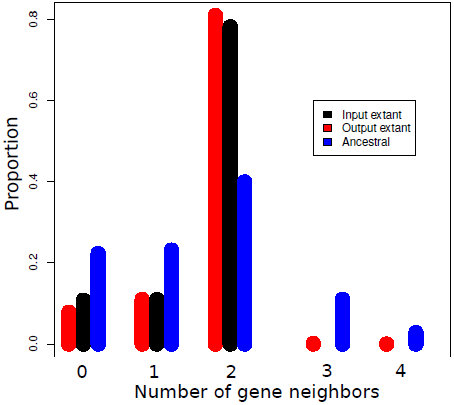
Number of syntenic neighbors of extant and ancestral genes. Distribution of the proportion of genes with a given number of neighbors in extant and ancestral genomes before and after adjacency prediction for the data set on 69 eukaryotes.

ART-DECO predicts 36,445 new extant adjacencies. As shown in Figure 3 (red bars) there is a significant increase in extant genes with 2 syntenic neighbors, as in a *bona fide* scaffolding, at the expense of a very small number of genes with more than two neighbors, corresponding to syntenic conflicts. Complementary computations show that more than 99.6% contigs in extant species remain linear (two genes having degree 1 and others degree 2), in spite of the large number of contig connections inferred in some species (*e.g.*, the number of contigs goes from 2,599 down to 1,864 for *Ailuropoda melanoleuca* or from 11,528 down to 7,930 for *Tarsius syrichta*). Figure 4 shows the percentage of improvement given by the method relatively to the initial data. Precisely, this percentage is obtained by computing 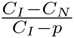 on extant species which are not completely assembled, where *C*_*I*_, resp. *C*_*N*_ and *p* are the number of contigs in the initial genome, resp. the number of contigs after adjacency inference by ART-DECO and the expected number of chromosomes. The figure shows that the more fragmented is the initial genome, the better ART-DECO improves it.

**Figure 4.**
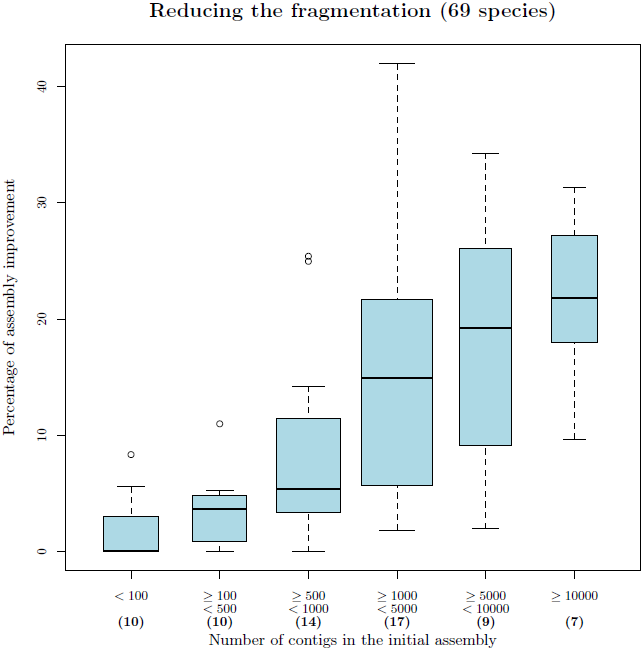
Percentage of improvement of genome assemblies, according to their initial fragmentation. Statistics are obtained for the 69 eukaryotes dataset, excluding genomes that are already well assembled (bold figures between parenthesis indicate cardinalities of classes).

Figure 5 shows the average degree of non-linearity of extant species with at least one non-linear contig, representing 43 of 69 species, computed only on non-linear contigs. Degree of non-linearity (*D*_*nl*_) correspond to supplementary degree of genes that are not consistent with a linear conformation and computed as follow:

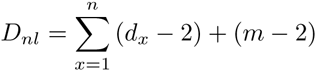

*n* = *Number of genes with degree >* 2
*d*_*x*_ = *Number of degree of gene x
m* = *Number of genes with degree* = 1

**Figure 5.**
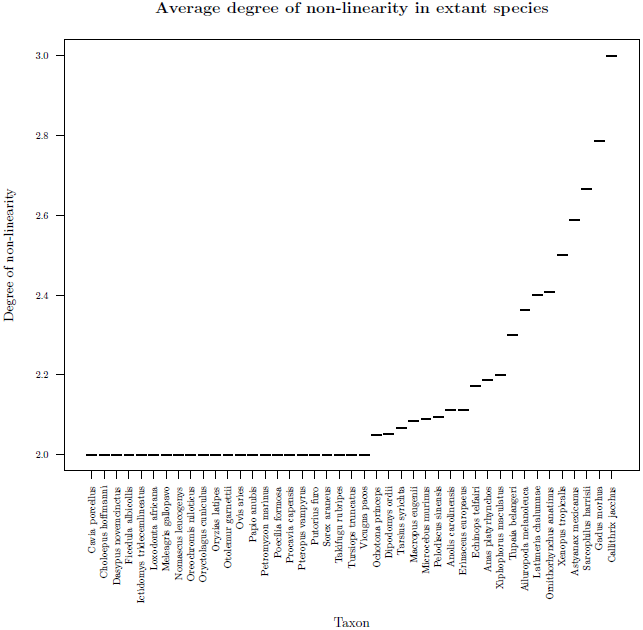
Distribution of average degree of non-linearity on non-linear contigs by extant species. On this graph, only species with at least one non-linear contig are shown, representing 43 of the 69 species. 23 of these species have an average degree of non-linearity of 2, meaning that their non-linear contigs contain an extra branch (one gene of degree 3 and one extra gene with degree 1).

On 43 species with non-linear contigs, 23 have non-linear contigs with only one extra branch. For the 20 remaining species, contigs are more branchy and few are circular.

We also reconstruct 1,547,546 ancestral adjacencies on 3,245,572 ancestral genes. As previously noted [33], errors in gene trees artificially inflate the number of ancestral genes computed with gene tree/species tree reconciliations. Nevertheless, the pattern of ancestral gene neighborhood shows mainly ancestral genes with 0, 1 or 2 neighbors, and some conflicts rapidly decreasing (Figure 3, blue bars). More than 92% inferred contigs in ancestral genomes are linear. Figure 6 presents the density histogram of average degree of non-linearity for ancestral species on inferred contigs that are not linear. As we can see most of the species have an average of degree non-linearity less than 20 meaning that in average non-linear ancestral contigs have a degree of non-linearity less than 20.Moreover, more than 50% of ancestral species have a degree of non-linearity less than 6 indicating that most of non-linear ancestral contigs are weakly branchy. However a large number of ancestral species have contigs strongly branchy and circular and need additional processes to obtain linear contigs. It is likely that better ancestral and extant genomes would result from better input gene trees.

**Figure 6.**
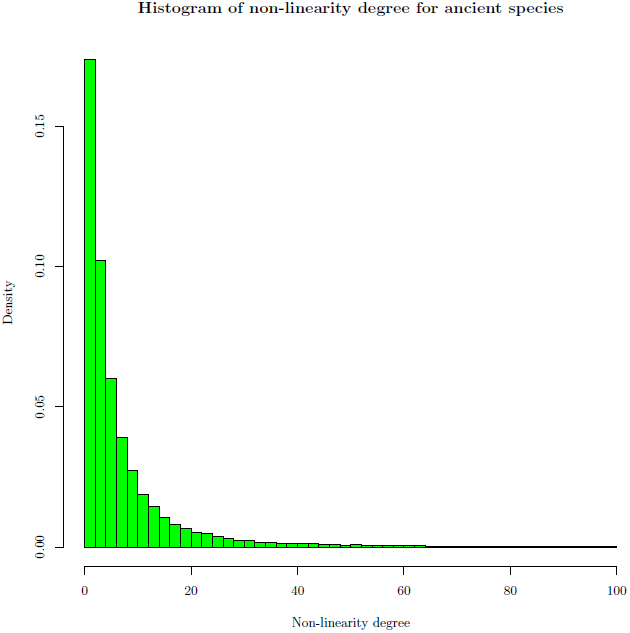
Density histogram of average degree of non-linearity for ancestral species on non-linear contigs. Most of the ancestral species have an average degree of non-linearity of 20 meaning that in average contigs of ancestral species reconstructed by ART-DECO have a degree of non-linearity less than 20. This figure shows that a large number are non-linear and additional operations are necessary to obtain linear contigs.

We analyze in details one predicted extant adjacency, in order to understand why it is present in the output of ART-DECO and not in the input. The adjacency we chose is randomly taken from the predicted ones between a contig and a chromosome in the panda genome (*Ailuropoda melanoleuca*). For this adjacency between genes RCSD1 and CREG1, we analyze gene neighborhoods around homologous adjacencies in others species. On Figure 7, we represent the species tree with information on evolutionary events that occurred along species tree (adjacency loss, duplication and gain, and gene loss and duplication) and adjacency status with color code on species name (red for species without RCSD1-CREG1 adjacency, blue for species with RCSD1-CREG1 adjacency in Ensembl database and green for species for which ART-DECO infers an adjacency between RCSD1 and CREG1 while not present in Ensembl). To illustrate the validity of new adjacencies inferred by ART-DECO we analyze the gene neighborhood around RCSD1-CREG1 adjacency. As, we can see on Figure 7, we observe that gene order and content is the same between cat (*Felis catus*), human (*Homo sapiens*) and panda except for ADCY10 gene (in black) that is not present in panda genome. The RCSD1-CREG1 adjacency is confirmed by adjacency support of *>* 99.96%, according to the exploration of the solution space. Due to the high fragmentation of panda genome and previous information it is reasonable to think that this adjacency is true. We observe the same results for the kangaroo rat (*Dipodomys ordii*) genome with gene content similarity with close species, high adjacency support (*>* 99.99%) and high genome fragmentation (9,720 contigs).

**Figure 7.**
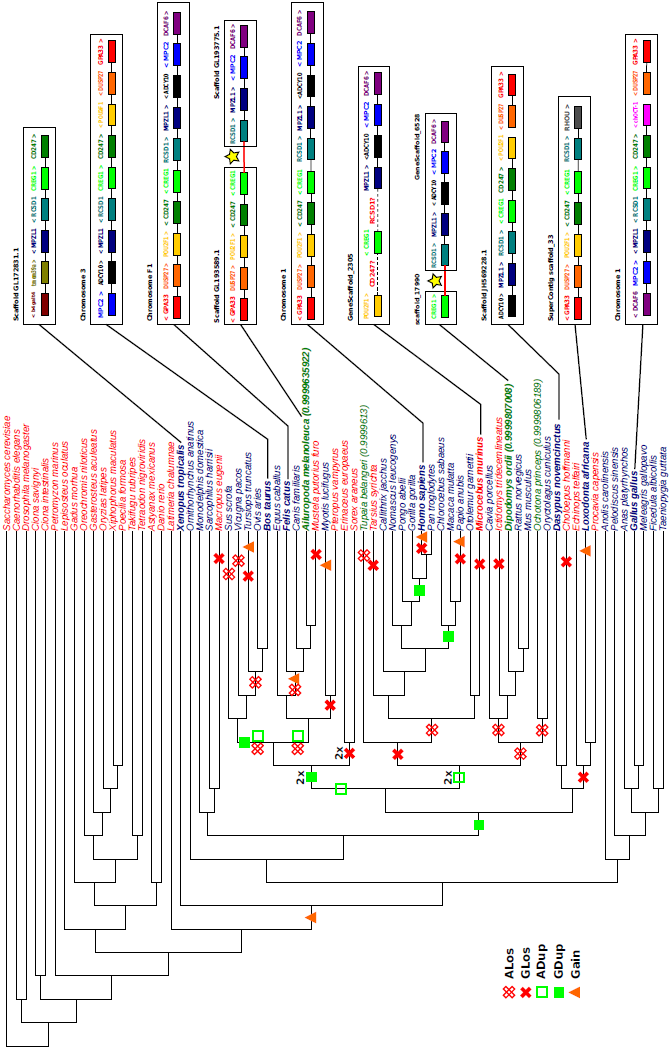
Evolutionary history of the adjacency between RCSD1 (turquoise) and CREG1 (light green). ART-DECO infers the creation of this adjacency at the root of Amniots. We integrated the different evolutionary events concerning the RCSD1-CREG1 adjacency along the species tree. Empty red crosses represents an adjacency losses (*i.e.*, cases where both adjacent genes are lost at the same time); each full red cross represents a gene loss (only one of the genes is lost); each empty green square indicates an adjacency duplication (places where the two adjacent genes are duplicated together); a full green square indicates a gene duplication and an orange triangle represents an adjacency gain. Color code for species name gives information on adjacency status. In red species, RCSD1 and CREG1 genes are not adjacent, while blue species host the RCSD1-CREG1 adjacency as described by Ensembl, and green species have the RCSD1-CREG1 adjacency inferred by ART-DECO (though it is absent from Ensembl). For green species, the adjacency support is indicated. For some species, representing most of the clades in Ensembl, we illustrate the gene content around the RCSD1-CREG1 adjacency, which illustrates strong similarities in the genomes of blue and green species.

This analysis also allowed to see a possible lack of data in Ensembl. As can be seen on the mouse lemur (*Microcebus murinus*) genome, there is no adjacency between CREG1 and RCSD1 because no RCSD1 gene has been annotated in Ensembl for this species. However, the gene content and order around CREG1 is very similar to that of close genomes (*e.g.*, human). Moreover, Ensembl contains an incomplete DNA sequence for the equivalent position of CD247 and RCSD1 genes in mouse lemur. This implies that the genes CD247 and RCSD1 could be present in mouse lemur but are not annotated.

### 39 mammals - validity

We switched to a smaller dataset to measure the validity of the method, because the computing time don’t allow too many replicates in the entire database. We selected all protein coding gene families from the 39 eutherian mammal genomes stored in the Ensembl database [30].

ART-DECO proposes 1,056,418 ancestral adjacencies and 22,675 new adjacencies in extant genomes. A proportion of 95% of these adjacencies have a *>* 0.9 support,meaning that they are present in over 90% of parsimonious solutions, computed as described in [31] for a *kT* value equal to 0.1 (chosen to ensure that the probability distribution over the solution space is highly dominated by optimal solutions).

Figure 8A shows the shape of extant genomes through the number or cumulative support of adjacencies incident to one gene. The distribution for all genes is plotted for extant genomes in the input and in the output, taking support into account or not. The figure shows that the genomes scaffolded with ART-DECO host more genes having exactly two neighbors (highest peak in the figure, the input is in black and output is in red). Peaks are integer numbers: unweighted measures have all their values integer while weighted measures still have peaks at integers. Complementary computations show that more than 99.7% contigs in extant species are indeed linear.

**Figure 8.**
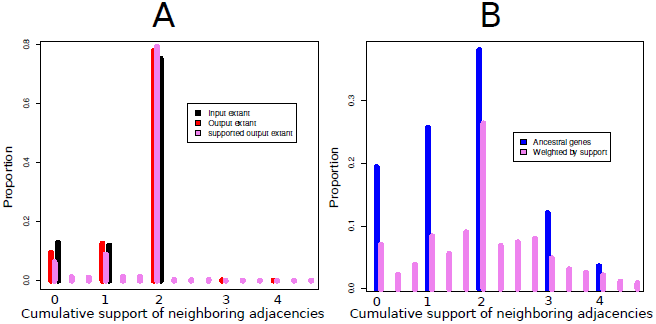
Weighted neighborhoods of extant (A) and ancestral genes (B). Distribution of the proportion of extant and ancestral genes with a given neighborhood weight in extant genomes before and after adjacency prediction, with or without support, for the data set on 39 mammals. The neighborhood weight of a gene is the sum of the supports of all adjacencies involving this gene. Continuous values are binned by intervals.

Figure 8B is the analog of Figure 8A but for ancestral genomes: blue for the number of neighbors in the version of ART-DECO without support (only one solution is given), and pink for the version of ART-DECO with support. The several peaks of the graph illustrate that ancestral genomes are not in the shape of disjoint paths, as we would expect it from linear genomes. This was already remarked in [12], and is likely due to errors in gene trees in the Ensembl database [34, 35]. Additional computations show that ancestral species indeed contain a larger proportion of nonlinear contigs: only 91.2% contigs are linear for those species, among which contigs hosting only one gene are more represented than in extant genomes. Thus, a small part of the inferred adjacencies are incorrect, leading to some artificially tree-like or cyclic contigs.

The bars with supports are more dispersed, as expected, because they take their values from non integer numbers. It puts the conflicts into perspective: when a gene has more than two neighbors, usually one adjacency is less supported.

We also performed experiments with artificial adjacency breaks as in the 7 tetrapods experiment. We removed from 1 to 75% input adjacencies from the human genome, and then from the horse genome. We chose the human and horse genome because of their phylogenetic position: one has many closely related genomes, while the other is rather distant from its closest neighbor inside the placentals. This allows us to measure the effect of the presence of closely related genomes in the given phylogeny. The two situations are very different because of the bias in taxonomic sampling around human. The presence of very close relatives in the data set makes the problem much easier for the human genome.

Indeed, as shown on Figure 9, the sensitivity (measured by the “*True positive*” rate as in the previous section, to keep a coherence and comparability with [29]) of the method is around 40% for the human genome, and 5% for the horse genome. The precision is high in all cases, decreasing with the number of broken adjacencies but the number of “*False positives*” stays quite low.

**Figure 9.**
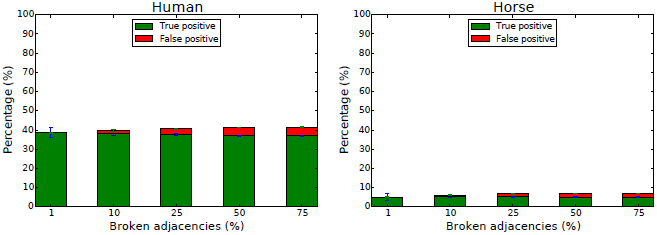
Capacity of ART-DECO to recover adjacencies after simulated breaks on human and horse genome.

The complexity of the data is a real issue here. While in a prepared, filtered data set of 7 tetrapods the sensitivity was above 80% in all cases, here with all genes from 39 genomes including duplications and losses, it is much lower in all cases.

From all data sets, we observe that the precision of ART-DECO is always high, while the sensitivity varies according to the conditions. So we can see the method as a rather sure predictor of a small number of scaffolding linkages, without the pretension to reconstruct fully assembled genomes.

## Discussion and Conclusion

Ancestral gene order reconstruction, when ancestral genes are given, can be seen as a scaffolding problem. Indeed ancestral genes may be seen as contigs, and finding an order between contigs is a similar problem in both extant and ancestral genomes. If this similarity had already been remarked and exploited in some way [24, 26, 36], a fully integrated approach has only recently been achieved by Aganezov *et al.* [29], with a method which was limited to universal unicopy genes and a small number of genomes. Extending DECO [12], a software aimed at reconstructing ancestral genomes and scaling up to dozens of genomes with possibly complex histories, we implement the additional possibility of scaffolding extant genomes in the same process, by handling equally ancestral and ancient genomes, with known and unknown parts in genome structures.

We demonstrate the efficiency of this approach on several eukaryote genomes data sets. It runs fast enough, proposes many additional supported adjacencies in extant genomes, and from several investigations we think we can state that such links are very likely to exist in reality. We are able to detect the less likely ones by assigning a support on ancestral and extant adjacencies by the same principle.

The main computational difference with the approach of Aganezov *et al.* [29] is that adjacencies are supposed to evolve independently. It has several notable consequences. The first one is the running time, because we switch from an NP-complete to a polynomial problem, and we are able to handle a large number of whole genomes. The second one is the shape of ancestral genomes. While methods modeling rearrangements [29] end up with *bona fide* genome structures, as linear arrangement of genes, our adjacency sets can be conflictual, both in ancestral and extant genomes. This means a gene can have more than two neighbors, unlike in real genomes. Whereas this can be seen as a serious drawback because genomes are not realistic, we would like to argue that it has several advantages, in addition to the running time. Indeed, the amount of conflicts can be a measure of uncertainty of the methods and data. It has been remarked many times that data sets, and in particular gene trees, are far from perfect. But better gene trees produce ancestral genomes with less conflicts [33]. Conflicts can point at problems that don’t necessarily concern the method itself, but give an overview of the quality of the data. This overview is lost if we force the data to fit in a linear structure. But if a linear ancestral genome is really needed, linearization techniques exist [37], even if we would argue for linearization techniques that also put into question the input data.

Some limitations would be still to overcome. For example we don’t handle the orientation of the genes. This would be desirable to have a finer account of ancestral and extant genomes, and to have a better fit between the *a priori* probability of an adjacency (computed in an oriented mode) and the reconstructed adjacencies. It would not be conceptually much complicated because adjacencies can be considered between gene extremities instead of between genes. But it would result in a loss of sensitivity because inversions of a single gene, which seem to be frequent, would fall into a rearrangement signal, increasing the probability to lose the traces of neighborhoods. We leave this open for a future work.

Another perspective is to be able to question extant adjacencies given in the input. In our framework they have probability 1, but a scaffolding is not necessarily only giving an order to the contigs. It can be inserting a contig inside another, or cutting a chimeric contig because a better arrangement can be proposed. Assembly errors are often numerous, not only because of a lack of information, but also because of false information [38]. It could be done by re-assigning an *a priori* probability to each extant adjacency, and not only to the ones outside the contigs. Finally, following the idea introduced in RACA [26], it could be interesting to pair the predictions of ART-DECO with sequence information such as mate-pairs or even physical or optical maps in order to integrate both evolutionary signal and sequencing data.

## Author’s contributions

YA, AC, CC, VB, ET and SB conceived the method and the tests. YA and SB implemented ART-DECO and YA tested it on all data sets. CC assigned the support scores on all retrieved adjacencies. YA, AC, CC, VB, ET and SB wrote the paper.

## Acknowledgements

This work is funded by the Agence Nationale pour la Recherche, Ancestrome project ANR-10-BINF-01-01. ART-DECO analyses benefited from the Montpellier Bioinformatics Biodiversity platform services. We thank Yann Ponty for technical help for patching DeClone to integrate the scaffolding and the exploration of the solution space.

## Additional Files

**Figure 10 -.**
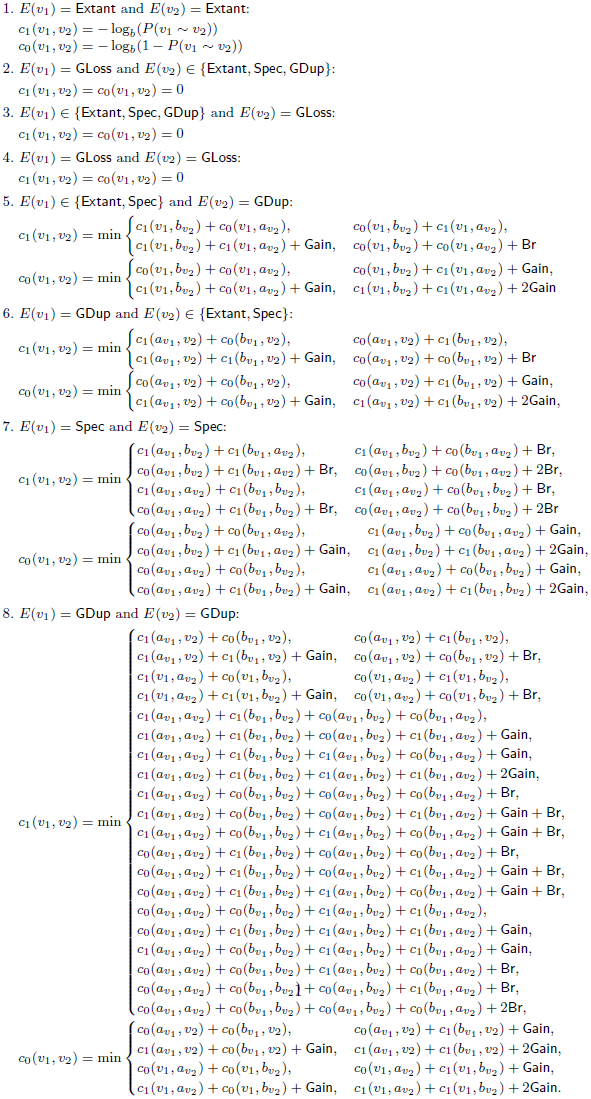
Additional file 1 ART-DECO dynamic programming scheme. Recurrence formulas used in ART-DECO to reconstruct adjacencies in ancestral and extant genomes.

**Figure 11 -.**
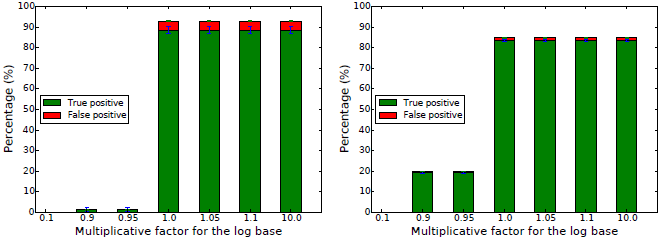
Additional file 2 Effect of number of adjacencies breaks on adjacencies recover in function of base log value. On the left, graph represents the number of adjacencies recovery in function of multiplicative factor for the log base for 50 simulated breaks in each species of 7 tetrapods dataset. On the right, the same graph but with 1050 simulated breaks for each species. As we can see the histogram profile is quietly similar between these two experiments and the one with 550 simulated breaks (see Figure 2). In conclusion, Number of adjacencies breaks didn’t impact the optimal value for the log base. (Values are available in Table 1).

## References

1. Raphael, B.J., Volik, S., Collins, C., Pevzner, P.A.: Reconstructing tumor genome architectures. Bioinformatics 19(Suppl. 2) (2003)

2. Fischer, A., Vázquez-Garcίa, I., Illingworth, C.J.R., Mustonen, V.: High-definition reconstruction of clonal composition in cancer. Cell Reports 7(5), 1740–1752 (2014)

3. McPherson, A., Roth, A., Ha, G., Shah, S.P., Chauve, C., Sahinalp, S.C.: Joint inference of genome structure and content in heterogeneous tumor samples. In: Research in Computational Molecular Biology. Lecture Notes in Computer Science, vol. 9029, pp. 256–258 (2015)

4. Hurst, L., Pál, C., Lercher, M.: The evolutionary dynamics of eukaryotic gene order. Nat Rev Genet 5(4), 299–310 (2004)

5. Swenson, K., Arndt, W., Tang, J., Moret, B.: Phylogenetic reconstruction from complete gene orders of whole genomes. In: Proceedings of the 6th Asia Pacific Bioinformatics Conference, pp. 241–250 (2008)

6. Sankoff, D.: Mechanisms of genome evolution: models and inference. Bulletin of the International Statistical Institute 47, 461–475 (1989)

7. Ma, J., Ratan, A., Raney, B.J., Suh, B.B., Zhang, L., Miller, W., Haussler, D.: DUPCAR: Reconstructing Contiguous Ancestral Regions with Duplications. Journal of Computational Biology 15(8), 1007–1027 (2008)

8. Chauve, C., Tannier, E.: A methodological framework for the reconstruction of contiguous regions of ancestral genomes and its application to mammalian genomes. PLoS Computational Biology 4(11), 1000234 (2008)

9. Alekseyev, M.A., Pevzner, P.A.: Breakpoint graphs and ancestral genome reconstructions. Genome Research 19(5), 943–957 (2009)

10. Ma, J.: A probabilistic framework for inferring ancestral genomic orders. In: IEEE International Conference on Bioinformatics and Biomedicine, BIBM, pp. 179–184 (2010)

11. Zheng, C., Sankoff, D.: On the PATHGROUPS approach to rapid small phylogeny. BMC Bioinformatics 12(Suppl. 1), 4 (2011)

12. Bérard, S., Gallien, C., Boussau, B., Szöllosi, G.J., Daubin, V., Tannier, E.: Evolution of gene neighborhoods within reconciled phylogenies. Bioinformatics 28(18), 382–388 (2012)

13. Hu, F., Lin, Y., Tang, J.: MLGO: phylogeny reconstruction and ancestral inference from gene-order data. BMC Bioinformatics 15, 354–359 (2014)

14. Reddy, T.B.K., Thomas, A.D., Stamatis, D., Bertsch, J., Isbandi, M., Jansson, J., Mallajosyula, J., Pagani, I., Lobos, E.A., Kyrpides, N.C.: The Genomes OnLine Database (GOLD) v.5: a metadata management system based on a four level (meta)genome project classification. Nucleic Acids Research 43(D1), 1099–1106 (2014). https://gold.jgi-psf.org/distribution

15. Simpson, J.T., Wong, K., Jackman, S.D., Schein, J.E., Jones, S.J.M., Birol, I.: ABySS: A parallel assembler for short read sequence data. Genome Research 19(6), 1117–1123 (2009)

16. Koren, S., Treangen, T.J., Pop, M.: Bambus 2: Scaffolding metagenomes. Bioinformatics 27(21), 2964–2971 (2011)

17. Salmela, L., Mäkinen, V., Välimäki, N., Ylinen, J., Ukkonen, E.: Fast scaffolding with small independent mixed integer programs. Bioinformatics 27, 3259–3265 (2011)

18. Gao, S., Sung, W.-K., Nagarajan, N.: Opera: Reconstructing optimal genomic scaffolds with high-throughput paired-end sequences. Journal of Computational Biology 18(11), 1681–1691 (2011)

19. Boetzer, M., Henkel, C.V., Jansen, H.J., Butler, D., Pirovano, W.: Scaffolding pre-assembled contigs using SSPACE. Bioinformatics 27(4), 578–579 (2011)

20. Gritsenko, A.A., Nijkamp, J.F., Reinders, M.J.T., de Ridder, D.: GRASS: A generic algorithm for scaffolding next-generation sequencing assemblies. Bioinformatics 28(11), 1429–1437 (2012)

21. Simpson, J.T., Durbin, R.: Efficient de novo assembly of large genomes using compressed data structures. Genome Research 22(3), 549–556 (2012)

22. Luo, R., Liu, B., Xie, Y., Li, Z., Huang, W., Yuan, J., He, G., Chen, Y., Pan, Q., Liu, Y., Tang, J., Wu, G., Zhang, H., Shi, Y., Liu, Y., Yu, C., Wang, B., Lu, Y., Han, C., Cheung, D.W., Yiu, S.-M., Peng, S., Xiaoqian, Z., Liu, G., Liao, X., Li, Y., Yang, H., Wang, J., Lam, T.-W., Wang, J.: SOAPdenovo2: an empirically improved memory-efficient short-read de novo assembler. GigaScience 1(1), 18 (2012)

23. Hunt, M., Newbold, C., Berriman, M., Otto, T.D.: A comprehensive evaluation of assembly scaffolding tools. Genome Biology 15(3), 42 (2014)

24. Husemann, P., Stoye, J.: Phylogenetic comparative assembly. Algorithms for Molecular Biology 5(1), 3–14 (2010)

25. Rajaraman, A., Tannier, E., Chauve, C.: FPSAC: Fast Phylogenetic Scaffolding of Ancient Contigs. Bioinformatics 29(23), 2987–2994 (2013)

26. Kim, J., Larkin, D.M., Cai, Q., Asan, Zhang, Y., Ge, R.-L., Auvil, L., Capitanu, B., Zhang, G., Lewin, H.A., Ma, J.: Reference-assisted chromosome assembly. Proceedings of the National Academy of Sciences (PNAS) 110(5), 1785–1790 (2013)

27. Kolmogorov, M., Raney, B., Paten, B., Pham, S.: Ragout - A reference-assisted assembly tool for bacterial genomes. Bioinformatics 30(12), 302–309 (2014)

28. Lin, Y., Nurk, S., Pevzner, P.A.: What is the difference between the breakpoint graph and the de Bruijn graph? BMC Genomics 15(Suppl. 6), 6 (2014)

29. Aganezov, S., Sitdykovaa, N., Alekseyev, M.A., AGCConsortium: Scaffold assembly based on genome rearrangement analysis. Computational Biology and Chemistry 57, 46–53 (2015)

30. Cunningham, F., Amode, M.R., Barrell, D., Beal, K., Billis, K., Brent, S., Carvalho-Silva, D., Clapham, P., Coates, G., Fitzgerald, S., Gil, L., Girón, C.G., Gordon, L., Hourlier, T., Hunt, S.E., Janacek, S.H., Johnson, N., Juettemann, T., Kähäri, A.K., Keenan, S., Martin, F.J., Maurel, T., McLaren, W., Murphy, D.N., Nag, R., Overduin, B., Parker, A., Patricio, M., Perry, E., Pignatelli, M., Riat, H.S., Sheppard, D., Taylor, K., Thormann, A., Vullo, A., Wilder, S.P., Zadissa, A., Aken, B.L., Birney, E., Harrow, J., Kinsella, R., Muffato, M., Ruffier, M., Searle, S.M.J., Spudich, G., Trevanion, S.J., Yates, A., Zerbino, D.R., Flicek, P.: Ensembl 2015. Nucleic Acids Research 43, 662–669 (2015)

31. Zanetti, J.P.P., Ponty, Y., Chauve, C.: Evolution of genes neighborhood within reconciled phylogenies: an ensemble approach. BMC Bioinformatics (2015). Brazilian Symposium on Bioinformatics 2014 special issue, to appear

32. Kasprzyk, A.: BioMart: Driving a paradigm change in biological data management. Database 2011, 049 (2011)

33. Boussau, B., Szollosi, G.J., Duret, L., Gouy, M., Tannier., E., Daubin, V.: Genome-scale coestimation of species and gene trees. Genome Research 23, 323–330 (2013)

34. Nouhati, E., Semeria, M., Lafond, M., Seguin, J., Boussau, B., Guéguen, L., El-Mabrouk, N., Tannier, E.: Genome evolution aware gene trees. https://hal.archives-ouvertes.fr/hal-01162963 (2015)

35. Rajaraman, A., Chauve, C., Ponty, Y.: Assessing the robustness of parsimonious predictions for gene neighborhoods from reconciled phylogenies. Lecture Notes in Computer Science, vol. 9096, pp. 260–271 (2015)

36. Luhmann, N., Chauve, C., Stoye, J., Wittler, R.: Scaffolding of ancient contigs and ancestral reconstruction in a phylogenetic framework. In: Proceedings of Brazilian Symposium on Bioinformatics. Lecture Notes in Computer Science, vol. 8826, pp. 135–143 (2014)

37. Manňuch, J., Patterson, M., Wittler, R., Chauve, C., Tannier, E.: Linearization of ancestral multichromosomal genomes. BMC Bioinformatics 13(Suppl. 19), 11 (2012)

38. Denton, J.F., Lugo-Martinez, J., Tucker, A.E., Schrider, D.R., Warren, W.C., Hahn, M.W.: Extensive error in the number of genes inferred from draft genome assemblies. PLoS Computational Biology 10(2), 1003998 (2014)

